# The structural OFF and ON states of myosin can be decoupled from the biochemical super-relaxed and disordered-relaxed states

**DOI:** 10.1101/2023.10.18.562891

**Authors:** Weikang Ma, Vivek P. Jani, Taejeong Song, Chengqian Gao, Henry Gong, Sakthivel Sadayappan, David A. Kass, Thomas C. Irving

## Abstract

There is a growing awareness that both thick filament and classical thin filament regulation play central roles in modulating muscle contraction. Myosin ATPase assays have demonstrated that under relaxed conditions, myosin may reside in either a high energy-consuming disordered-relaxed (DRX) state available for binding actin to generate force, or in an energy-sparing super-relaxed (SRX) state unavailable for actin binding. X-ray diffraction studies have shown the majority of myosin heads are in a quasi-helically ordered OFF state in a resting muscle and that this helical ordering is lost when myosin heads are turned ON for contraction. It has been assumed that myosin heads in SRX and DRX states are equivalent to the OFF and ON state respectively and the terms have been used interchangeably. Here, we use X-ray diffraction and ATP turnover assays to track the structural and biochemical transitions of myosin heads respectively induced with either omecamtiv mecarbil (OM) or piperine in relaxed porcine myocardium. We find that while OM and piperine induce dramatic shifts of myosin heads from the OFF to ON states, there are no appreciable changes in the population of myosin heads in the SRX and DRX states in both unloaded and loaded preparations. Our results show that biochemically defined SRX and DRX can be decoupled from structurally-defined OFF and ON states. In summary, while SRX/DRX and OFF/ON transitions can be correlated in some cases, these two phenomena are measured using different approaches, do not necessarily reflect the same properties of the thick filament and should be investigated and interpreted separately.

**Significance:** Myosin based thick filament regulation is now known to be critical for muscle contraction with myosin dysregulation found in hypertrophic and dilated cardiomyopathies. While previously thought to be synonymous, this study finds that biochemical and structural thick filament disengagement are distinct properties and should be investigated as independent phenomena. Understanding the details of thick filament regulation will be of great relevance to defining sarcomere-level dysfunction in myopathies and understanding and better designing and testing sarcomere therapies aimed at reversing them for treatment of cardiomyopathy.

## Introduction

Regulation of vertebrate striated muscle contraction has been regarded as a calcium (Ca^2+^) mediated thin filament-based mechanism. Upon excitation signaling, Ca^2+^ enters the cytosol to bind to troponin-C on the thin filament, triggering a series of conformational changes to displace tropomyosin from myosin-binding sites on actin, allowing for actin-myosin crossbridge formation and thus force generation. Initial binding of myosin to the unblocked sites results in a full cooperative activation of the thin filament to augment force (1, 2). This classical Ca^2+^-mediated thin filament-based regulation mechanism assumes that all myosin heads are free to bind actin once the actin binding sites are available. However, this picture appears to be incomplete, and we now realize that muscle regulation requires both thick and thin filament based mechanisms to fully activate the sarcomere (3).

Thick filament based regulation in vertebrate muscle was first bought to our attention when Roger Cooke and colleagues discovered that under resting conditions, myosin can exist either in a disordered-relaxed (DRX) state with a higher ATP consumption rate (∼ 0.03 s^−1^) or in an energy-sparing, low ATP consumption (∼0.003 s^−1^) state, known as the super-relaxed (SRX) state (4, 5). SRX and DRX are, strictly speaking, biochemically defined terms that depend on ATP consumption rates of myosin heads. Subsequent studies suggested that myosin heads in the SRX state might be sequestered on the surface of thick filament making them unavailable for binding to actin, whereas heads in the DRX state are free to bind to actin and generate force (6-9). The relative proportions of myosin heads in SRX and DRX states under resting conditions have been proposed to be related to the amount of force produced and to be largely responsible for the hypo- and hyper-contractility observed with hereditary cardiomyopathies and more recently, end-stage heart failure (9-12). This concept motivated the development of direct myosin interventions as a therapeutic strategy to correct contractile abnormalities in myopathies culminating in the FDA approval of mavacamten (Camzyos) to treat obstructive HCM (13, 14)

Another aspect of thick filament activation was brought to the forefront by Linari and colleagues, who proposed a mechanosensing-based thick filament activation model (15). In the resting state, the majority of the myosin heads are quasi-helically ordered on the surface of the thick filament backbone. These myosin heads, defined to be in the OFF state, produce the characteristic myosin-based layer line reflections in X-ray fiber diffraction patterns (16-18). The helical ordering is lost when myosin heads are turned ON to participate in contraction(15, 19, 20). In the mechano-sensing model, once the thin filament is turned on by influx of Ca^2+^, a small portion of constitutively ON heads, assumed to be constantly searching for binding sites on actin, will bind to actin and generate small amounts of force that strain the thick filament. This strain then results in converting more myosin heads from the OFF state to the ON state, a behavior lost in end-stage heart failure (3, 15).

It has been generally assumed that myosin heads in the biochemically-defined SRX and DRX state are equivalent to the structurally-defined OFF and ON state respectively (3, 6, 21), so that the terms SRX state and OFF state are often used interchangeably. Here we used X-ray diffraction and ATP turnover assays to track the structural and biochemical transitions induced by either omecamtiv mecarbil (OM) or piperine respectively under resting conditions. OM and piperine were chosen in this study as OM was the first myosin activator to undergo, ultimately unsuccessful, clinical trials as a myosin activator (22), and piperine, a phenol derivative of black pepper, that has been shown to change the portion of myosin heads in the SRX state in fast skeletal muscle but not in slow skeletal muscle and cardiac muscle (23, 24). We show that while OM and piperine induces dramatic shifts of myosin heads from the OFF to the ON states in skinned porcine cardiac muscle, there are no appreciable changes in the populations of myosin heads in the SRX and DRX states. These results reveal that the biochemically-defined SRX and DRX states can be decoupled from the structurally-defined OFF and ON states, indicating that these behaviors do not necessarily reflect the same underlying phenomena.

## Results

### Structural changes of permeabilized porcine myocardium with OM and piperine

We studied X-ray diffraction patterns obtained from relaxed permeabilized porcine myocardium at different concentrations of OM and piperine under resting conditions (pCa8). Qualitatively, permeabilized porcine myocardium showed characteristic relaxed myosin-based layer lines (MLL1 and MLL2) in the absence of myosin activators, that diminished in intensity with increasing activator concentration, until they were no longer visible in the presence of the highest concentrations of OM (∼10 *μ*M) and piperine (∼50 *μ*M) used in this study (Fig 1A and 1B, respectively). The equatorial intensity ratio (I_1,1_/I_1,0_), an indicator of the proximity of myosin heads to actin in relaxed muscle (16), increased monotonically as a function of either OM (Fig 1C, Table S1) or piperine (Fig 1D, Table S2) concentrations, indicating a shift of myosin heads away from the thick-filament backbone towards the thin filament at increased concentration of each activator.

**Figure 1.**
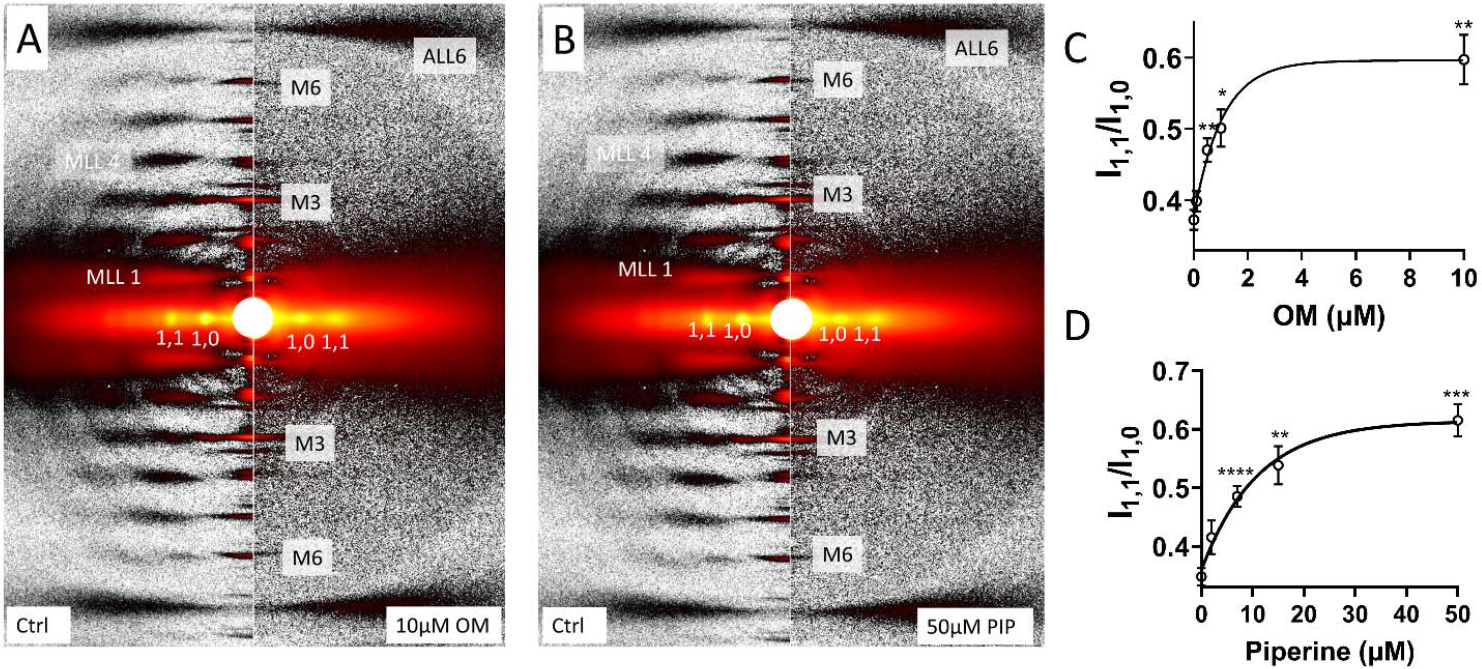
X-ray diffraction patterns from permeabilized porcine myocardium in relaxing solution in the absence and presence of myosin activators. (**A**), X-ray diffraction patterns from relaxed muscle in the absence (left panel) and presence (right panel) of 10 µM omecamtiv mecarbil (OM). (**B**), X-ray diffraction patterns from relaxed muscle in the absence (left panel) and presence (right panel) of 50 µM piperine (PIP). Equatorial intensity ratio (I_1,1_/I_1,0_) at different concentration of omecamtiv mecarbil (OM) **(C)** and piperine **(D)**. Myosin heads moves radially closer to actin as OM and piperine concentration increases. The results are given as mean ± SEM with *p* values were calculated from RM one-way ANOVA with Dunnett’s multiple comparisons test compare to control (0 µM activator). Symbols on figures: *: p<0.05, **: p<0.01, ***: p<0.001,****: p<0.0001.

Both I_MLL1_ and I_M3_ decreased slightly with 0.1 µM OM treatment, and further decreased monotonically (up to 70% at 10 µM OM) at higher concentrations (Fig 2A & 2C, Table 1). Similarly, both I_MLL1_ and I_M3_ decreased slightly at 2 µM piperine and further decreased monotonically at higher concentrations (Fig 2B & 2D, Table 1). The intensity of the sixth order myosin-based meridional reflection (M6) arises primarily from the thick filament backbone. The spacing of M6 reflection (S_M6_), reflecting the thick filament backbone periodicity (16), increases as the concentration of either OM and piperine increases (Fig 2E & 2F, Table 1). Both an increase in the thick filament backbone periodicity, indicated by increased S_M6_, and a reduction in the degree of the helical ordering of the myosin heads, indicated by reduced I_MLL1_ and I_M3_, are characteristic signatures of the structurally-defined OFF to ON transition of myosin (3, 16, 25). Taken together, these findings show both OM and piperine result in substantial disruption of the helical ordering of myosin heads on the surface of the thick filament backbone resulting in release of myosin heads to move closer to actin; i.e. an OFF to ON transition.

**Table 1.** X-ray diffraction pattern changes in the presence of increasing concentration of omecamtiv mecarbil (OM) and piperine (PIP)

| | Interpretation | 0 μM OM | 0.1 μM OM | 0.5 μM OM | 1 μM OM | 10 μM OM | Half-value |
| --- | --- | --- | --- | --- | --- | --- | --- |
| $I_{1,1}/I_{1,0}$ | Proximity of myosin heads to actin | $0.37 \pm 0.014$ | $0.40 \pm 0.014$ | $0.47 \pm 0.016^{**}$ | $0.50 \pm 0.026^*$ | $0.60 \pm 0.035^{**}$ | 0.74 μM |
| $I_{MLL1}$ (a.u.) | Helical ordering of the myosin heads | $10.76 \pm 0.71$ | $8.36 \pm 0.84$ | $6.88 \pm 0.60^{**}$ | $6.16 \pm 0.65^*$ | $3.72 \pm 0.26^{***}$ | 0.56 μM |
| $I_{M3}$ (a.u.) | Axial ordering of the myosin heads | $6.67 \pm 0.50$ | $5.52 \pm 0.44$ | $4.68 \pm 0.35^{**}$ | $3.87 \pm 0.19^{****}$ | $3.30 \pm 0.16^{****}$ | 0.41 μM |
| $S_{M6}$ (nm) | Thick filament backbone periodicity | $7.186 \pm 0.002$ | $7.192 \pm 0.002$ | $7.199 \pm 0.002^{***}$ | $7.201 \pm 0.002^{**}$ | $7.207 \pm 0.002^{****}$ | 0.47 μM |

pattern changes in the presence of increasing concentration of omecamtiv mecarbil (OM) and piperine (PIP)
| | Interpretation | 0 μM PIP | 2 μM PIP | 7 μM PIP | 15 μM PIP | 50 μM PIP | Half-value |
| --- | --- | --- | --- | --- | --- | --- | --- |
| $I_{1,1}/I_{1,0}$ | Proximity of myosin heads to actin | $0.35 \pm 0.015$ | $0.42 \pm 0.029$ | $0.49 \pm 0.018^{****}$ | $0.54 \pm 0.033^{**}$ | $0.62 \pm 0.028^{**}$ | 1.68 μM |
| $I_{MLL1}$ (a.u.) | Helical ordering of the myosin heads | $14.09 \pm 1.36$ | $9.55 \pm 1.02$ | $7.21 \pm 0.88^{***}$ | $5.34 \pm 0.60^{**}$ | $3.90 \pm 0.54^{***}$ | 3.36 μM |
| $I_{M3}$ (a.u.) | Axial ordering of the myosin heads | $10.12 \pm 1.19$ | $7.19 \pm 1.02$ | $5.24 \pm 0.41^{**}$ | $4.25 \pm 0.51^{**}$ | $3.13 \pm 0.35^{**}$ | 3.29 μM |
| $S_{M6}$ (nm) | Thick filament backbone periodicity | $7.178 \pm 0.004$ | $7.185 \pm 0.003$ | $7.191 \pm 0.003^{**}$ | $7.197 \pm 0.002^{**}$ | $7.200 \pm 0.002^{**}$ | 5.23 μM |

Values are given as mean ± SEM. The $p$ values were calculated from RM one-way ANOVA with Dunnett's multiple comparisons test compare to control (0 μM OM or piperine). The half-value was calculated by fitting the values of each reflection in a function of activator concentration to an exponential function. n=8. Symbols on table: \*: $p < 0.05$ , \*\*: $p < 0.01$ , \*\*\*: $p < 0.001$ , \*\*\*\*: $p < 0.0001$ .

**Figure 2.**
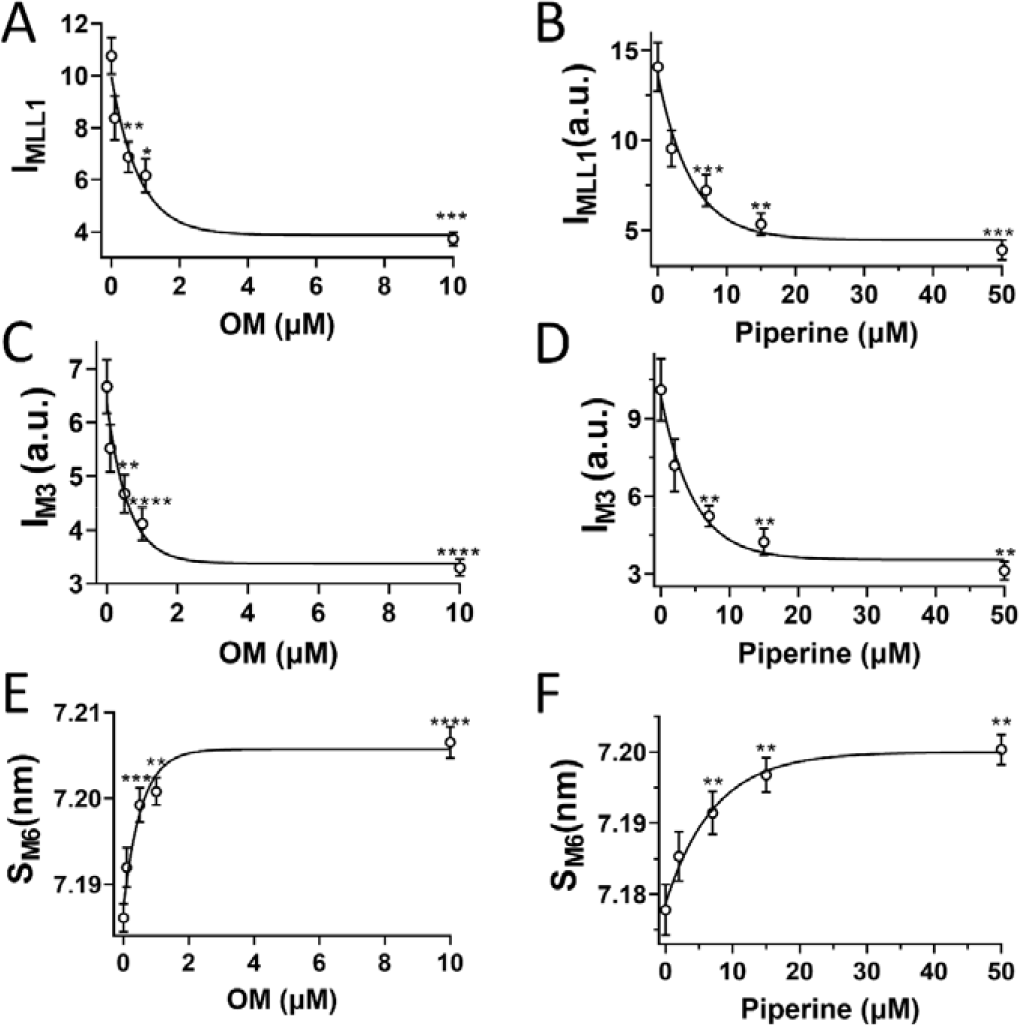
Thick filament structural changes in the presence of myosin activators. The intensity of the first order myosin-based layer line (I_MLL1_) in different concentrations of OM (**A**) and piperine (**B**). The third order myosin based meridional reflection (I_M3_) in different concentrations of OM (**C**) and piperine (**D**).The spacing of the sixth order myosin based meridional reflection (S_M6_) in different concentrations of OM (**E**) and piperine (**F**) Myosin heads moves from the helically ordered OFF states to disordered ON states as activators concentration increase. Results are given as mean ± SEM with *p* values were calculated from RM one-way ANOVA with Dunnett’s multiple comparisons test compare to control. Symbols on figures: *: p<0.05, **: p<0.01, ***: p<0.001,****: p<0.0001.

### Changes in ATP turnover rate in permeabilized porcine myocardium with OM and piperine

To test the hypothesis that structurally-defined OFF/ON states myosin heads are strongly correlated to biochemically-defined SRX/DRX states, we next examined the proportion of myosin heads in SRX and DRX states with and without both activators. OM (5 μM) or piperine (50 μM) was applied to permeabilized myocardial strips under unloaded conditions, and the decay rate of fluorescent MANT-ATP provided a measure of ATP turnover rate. The relative proportions of the myosin heads in the DRX and SRX states and the time constant of the fast phase (T1) and the slow phase (T2) were calculated by fitting the fluorescence decay profile with a 2-phase exponential decay function (c.f. methods). We observed overlapping fluorescence decay signals in control (un-treated) and OM/piperine treated tissues. Neither the fraction of the myosin heads in the SRX state nor the T1 and T2 values change significantly in the presence of either OM or piperine (Fig 3, Table 2).

**Table 2.** ATP turnover assay before and after either omecamtiv mecarbil (OM) or piperine (PIP) treatment.

|  | Unloaded myocardium bundles |  |  |  | Loaded cardiomyocytes |  |  |  |  |  |
| --- | --- | --- | --- | --- | --- | --- | --- | --- | --- | --- |
| | Ctrl (n=15) | OM(n=11) | PIP (n=9) | $p$ values | Ctrl (n=10) | OM (n=10) | $p$ values | Ctrl (n=10) | PIP (n=10) | $p$ values |
| %SRX | $49.1 \pm 6.9$ | $58.5 \pm 7.1$ | $42.32 \pm 9.1$ | ns | $47.71 \pm 2.15$ | $43.62 \pm 2.75$ | ns | $42.99 \pm 4.32$ | $38.57 \pm 3.20$ | ns |
| T1 (s) | $24.20 \pm 1.67$ | $25.52 \pm 1.99$ | $27.65 \pm 2.67$ | ns | $8.74 \pm 0.82$ | $5.9 \pm 0.78$ | <0.01 | $9.20 \pm 1.68$ | $7.24 \pm 0.60$ | ns |
| T2 (s) | $129.6 \pm 13.0$ | $114.5 \pm 14.1$ | $145.9 \pm 20.3$ | ns | $271.5 \pm 27.9$ | $213.4 \pm 41.6$ | ns | $321.2 \pm 61.3$ | $270.9 \pm 30.82$ | ns |

**Figure 3.**
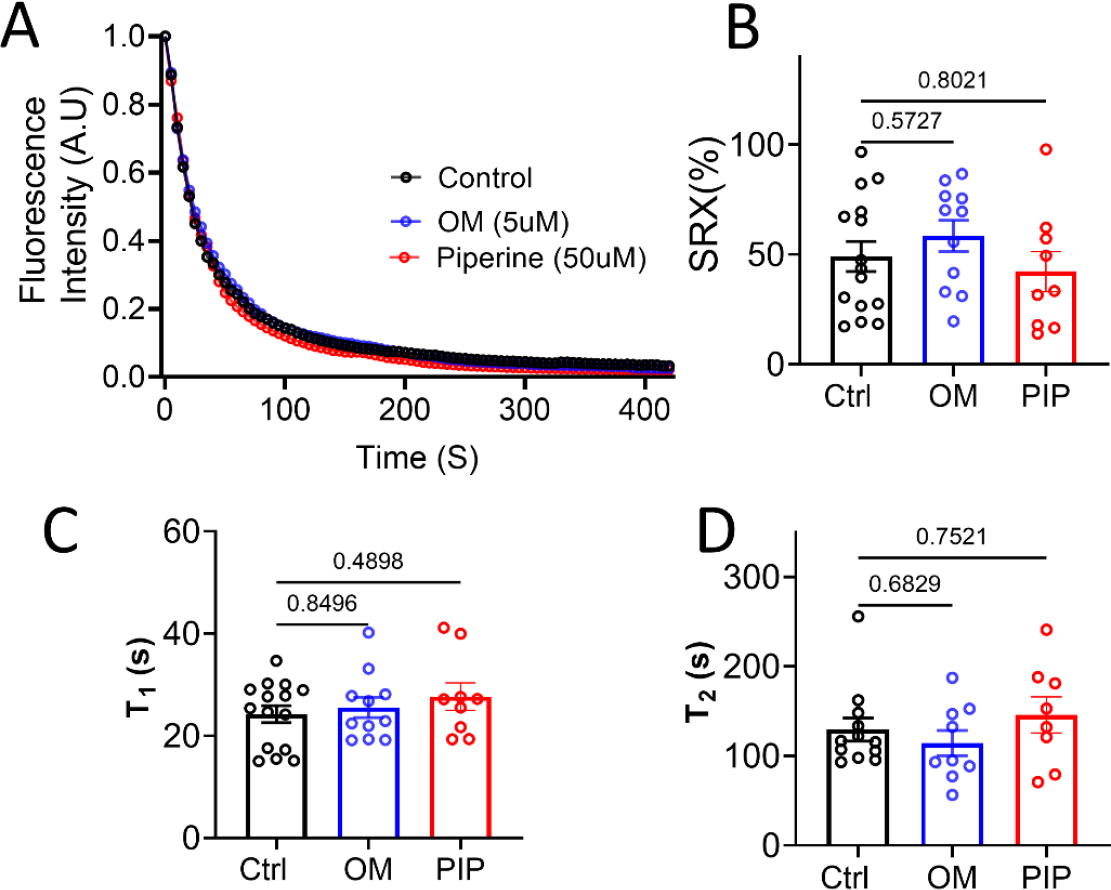
ATP turnover assays of unloaded permeabilized porcine myocardium bundles treated with OM and piperine (PIP). **A:** Mant-ATP dissociation over time curves in control (black), 5 μM OM (blue) and 50 μM piperine (red). **B:** The percentage of myosin heads in the SRX state (% SRX) with and without OM and piperine. **C**: The time constant of the fast phase (T_1_) with and without OM and piperine treatment. **D**: The time constant of the slow phase (T_2_) with and without the OM and piperine treatment. The results are given as mean ± SEM, with *p* values as shown in the figure calculated from unpaired Brown-Forsythe and Welch ANOVA test.

While unloaded assays are commonly used to assess SRX/DRX transitions, it has now been shown that the population of myosin heads in SRX vs DRX is sensitive to the sarcomere length in both porcine (28) and human myocardium (32). Sarcomere length is not controlled for in unloaded assays, contributing to increased variance in the measurements. Furthermore, the diffusion of free nucleotides out of relatively large multicellular muscle bundles takes 10-15 s, overlapping with the fast phase time constant (T1) in the ATP turnover assay, complicating data interpretation (7). To mitigate these potential limitations, we also used a recently developed loaded ATP turnover procedure (25) in permeabilized single cardiomyocytes (CMs) and measured the proportions of myosin heads in the SRX/DRX states in paired pre- and post-incubations with both compounds of interest. Similar to the results obtained from unloaded fiber bundles, we find no significant changes in the proportion of myosin heads in the SRX state (OM p=0.28; piperine p=0.43) before and after OM or piperine treatment (Fig 4 and Table 2). However, we observed a significant 35% reduction (Table 2) in T1 with 1 *μ*M OM (Figure 4E), suggesting that OM increases the ATPase activity of DRX myosin heads. In summary, in all cases, there were no significant changes in the fraction of the myosin heads in the SRX state despite the increase in equatorial intensity ratio previously observed.

**Figure 4.**
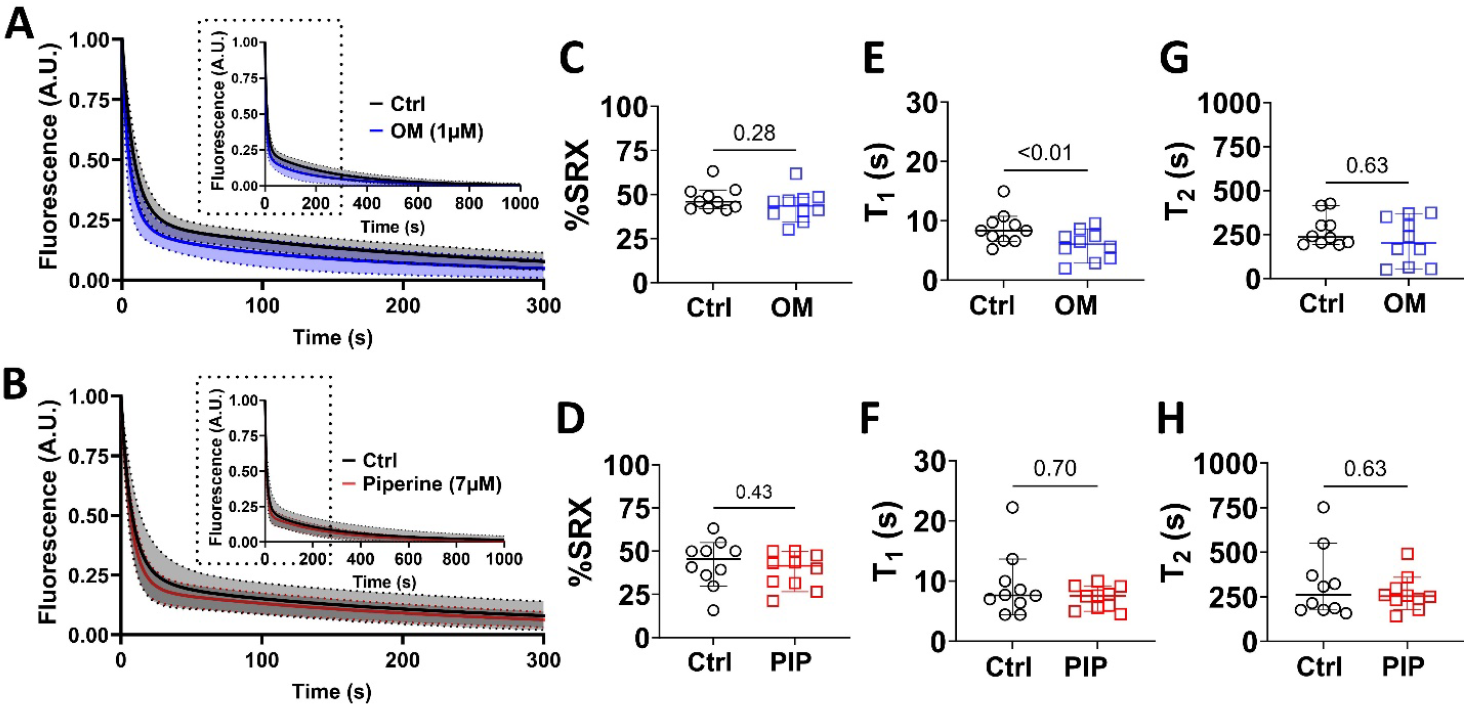
ATP turnover assays of loaded permeabilized porcine CMs before and after OM and piperine (PIP) treatment. **A:** Mant-ATP dissociation over time curves in control (black) and 1 μM OM (blue). **B:** Mant-ATP dissociation over time curves in control (black) and 7 μM piperine (red). The percentage of myosin heads in the SRX state (% SRX) before and after OM (**C**) and piperine (**D**) treatment. The time constant of the fast phase (T_1_) before and after the OM (**E**) and piperine (**F**) treatment. The time constant of the slow phase (T_2_) before and after the OM (**G**) and piperine (**H**) treatment. The results are given as mean ± SEM with *p* values as shown in the figure calculated from Wilcoxon matched-pairs signed rank t tests.

The results are given as mean ± SEM, with *p* values are calculated from unpaired Brown-Forsythe and Welch ANOVA test for unloaded myocardium bundles experiments and *p* values calculated from Wilcoxon matched-pairs signed rank t tests for Loaded cardiomyocytes experiments. Symbols on table, ns: not significant

## Discussion

It has been often observed that the structurally-defined OFF and ON states of myosin are correlated with the biochemically defined SRX and DRX states (3, 6, 10, 26) and, consequently, are widely assumed to reflect the same underlying phenomena. Mavacamten, a small molecule myosin inhibitor, treated myocardium is one case where transitions of myosin heads from the DRX to SRX are well correlated to ON to OFF transitions (7, 27). Additionally, it has been shown that, at least in one cohort of right ventricle heart failure patients, increases in the population of myosin heads in the biochemical SRX state and structural OFF state might be the underlying cause of depressed contractile force associated with right ventricular failure (28). Myosin activators that were able to recruit myosin from the OFF and the SRX states were therefore proposed to be one possible therapy for these patients. So far, only deoxy-ATP (29, 30) and EMD-57033 (25) have been shown, when used as tool compounds, to be able to recruit myosin from the OFF and SRX states to ON and DRX states, respectively.

The structural basis for the SRX has been proposed to be the interacting head motif (IHM), in which a pair of the myosin heads interact with each other and the S2 segment causes them to be held close to the thick filament backbone (6, 31, 32). While our definition of the helically ordered OFF state does not address the precise configuration, and possible heterogeneity, of the OFF-state myosin heads, a recent cryo-EM study on isolated human cardiac thick filaments showed that there are three distinct IHM configurations that are quasi-helically ordered on the surface of the thick filament in the presence of mavacamten (33). This suggests that a substantial fraction of the myosin heads in the structurally defined OFF state in resting muscle are likely to be in one of these IHM states. Here, our X-ray diffraction data showed that increased concentration of OM and piperine induce a dramatic reduction in the helical ordering of the myosin heads (Fig 2) accompanied with a radial movement of the myosin heads towards actin (Fig 1) at pCa 8. These data strongly indicate an OFF to ON transition of myosin heads induced by OM and piperine. ATP turnover assays using both unloaded porcine fiber bundles and loaded cardiomycytes, however, showed that OM and piperine did not significantly alter the relative portion of the myosin heads in the SRX and DRX states. These findings refute the notion that the biochemically defined SRX and DRX states are necessarily manifestations of the structurally-defined OFF and ON states, but rather that these behaviors are distinct. We conclude that a biochemical SRX to DRX transition of myosin does not necessarily imply a structural OFF to ON transition and that a structural OFF to ON transition does not necessarily imply a biochemical SRX to DRX transition. It is worth noting that suggestions of possible disconnects between the biochemical and structural states of myosin have previously appeared in the literature. In relaxed porcine myocardium, while structural OFF to ON and biochemical SRX to DRX transitions increased in response to dATP (29), DRX myosin heads increased linearly with increasing dATP concentrations, whereas the structural OFF to ON response was non-linear. Chu and colleagues showed that mavacamten, a small molecule myosin inhibitor, has a greater effect on the increase of population of myosin in the SRX state than the increase in number of myosin heads in the IHM state (31).

It is clear that the biochemically-defined SRX and DRX states can respond differently to physiological perturbations and experimental interventions from the structurally defined OFF and ON states and that they should not necessarily be considered to represent the same underlying phenomena. These findings also have direct translational significance. First of all, it is clear that for any compound that might target the SRX/DRX or the OFF/ON equilibrium as a therapeutic route for both heart and skeletal muscle diseases, one cannot assume they are necessarily coupled. Secondly, ongoing studies in our laboratories indicate that decoupling of biochemical SRX/DRX and structural OFF/ON transitions probably occurs more widely than presented here, and rather than a peculiarity of the specific compounds studied here, the demonstration that the two phenomenon do not act in lock-step is of broad physiological and pathological significance for health and disease. We suggest, therefore, that both SRX/DRX and structural OFF/ON state transition measurements should be done as part of a comprehensive study either of the mode of action of candidate therapeutic compounds, or underlying disease mechanisms, to better inform further decision-making in therapeutic strategies.

## Materials and Methods

### Muscle preparation for X-ray diffraction

Porcine myocardium was prepared as described previously (34, 35). Briefly, frozen porcine left ventricle wall was thawed in skinning solution (pCa8 solution: 91 mM K^+^-propionate, 3.5 mM MgCl, 0.16 mM CaCl_2_, 7 mM EGTA, 2.5 mM Na_2_ATP, 15 mM creatine phosphate, 20 mM Imidazole) plus 30 mM BDM, 1% Triton-X100 at room temperature for 1h before dissection into smaller strips. Myocardium strips were skinned at room temperature for 2 hours. The fiber bundles were further dissected into preparations with a length of 5 mm with a diameter of 200 µm prior to attachment of aluminum T-clips to both ends in pCa 8 solution with 3% dextran on ice for the day of the experiment. X-ray diffraction experiments were performed at the BioCAT beamline 18ID at the Advanced Photon Source, Argonne National Laboratory (36). The X-ray beam energy was set to 12 keV (0.1033 nm wavelength) and the incident flux was set to ∼5×10^12^ photons per second. The specimen to detector distance was about 3 m. The muscle was incubated in a customized chamber with one end attached to a force transducer (Model 402B Aurora Scientific Inc., Aurora, ON, Canada) and Kapton™ windows in the X-ray path in pCa 8 solution. The experiment was performed between 28 °C and 30 °C at a sarcomere length of 2.3 µm as adjusted by laser diffraction. X-ray diffraction patterns were collected on a MarCCD 165 detector (Rayonix Inc., Evanston IL) with a 1 s exposure time as a function of five increasing Omecamtiv mecarbil OM (0 µM, 0.1 µM, 0.5 µM, 1 µM, and 10 µM) and piperine (0 µM, 2 µM, 7 µM, 15 µM, and 50 µM) concentrations. OM was purchased from AdooQ Biosciences (Irvine, CA), and piperine from Millipore Sigma (St Louis, MO). To minimize radiation damage, the muscle was oscillated along its horizontal axes at a velocity of 1 mm/s and moved vertically after each exposure to avoid overlapping X-ray exposures. Two to three patterns were collected under each condition and the spacings and intensities of selected X-ray reflections extracted from these patterns were averaged.

### X-ray data analysis

The data were analyzed using the MuscleX software package developed at BioCAT (37). Briefly, the equatorial reflections were measured by the “Equator” routine in MuscleX as described previously (38). The intensities and spacings of meridional and layer line reflections were measured by the “Projection Traces” routine in MuscleX as described previously (27, 39). To compare reflection intensities under different conditions, the measured intensities of X-ray reflections were normalized to the intensities of the sixth-order actin-based layer line (29). The results are given as mean ± SEM with *p* values were calculated from RM one-way ANOVA with Dunnett’s multiple comparisons test compare to control (0 µM activator). Symbols on figures: ns: not significant, *: p<0.05, **: p<0.01, ***: p<0.001,****: p<0.0001. N and n indicate biological and technical repeats respectively.

### Unloaded ATP turnover assays in permeabilized porcine myocardium bundles

Permeabilized porcine cardiac tissue was prepared as described previously (40, 41). Briefly, previously frozen porcine cardiac tissue was cut in small pieces and skinned for six hours on ice in a skinning buffer (100 mM NaCl, 8 mM MgCl_2_, 5 mM EGTA, 5 mM K_2_HPO_4_, 5 mM KH_2_PO_4_, 3 mM NaN_3_, 5 mM ATP, 1 mM DTT, 20 mM BDM and 0.1% (v/v) Triton X-100 at pH 7.0). The skinning buffer was changed every two hours. After skinning, the permeabilized fibers were placed in a glycerinating buffer overnight at 4 °C. Then, samples were stored in fresh glycerinating solution (120 mM K acetate, 5 mM Mg acetate, 5 mM EGTA, 2.5 mM K_2_HPO_4_, 2.5 mM KH_2_PO_4_, 50 mM MOPS, 5 mM ATP, 20 mM BDM, 2 mM DTT, and 50% (v/v) glycerol at pH 6.8) at -20°C until use. To measure the SRX/DRX states of myosin, skinned cardiac tissue was further cut into a small bundle of fibers in cold glycerol buffer. Both sides of fiber was secured with double sided tape in a flow chamber (thickness ∼270 µm) as described previously (41). Samples were washed five times with rigor buffer (120 mM K acetate, 5 mM Mg acetate, 5 mM EGTA, 2.5 mM K_2_HPO_4_, 2.5 mM KH_2_PO_4_, 50 mM MOPS, and 2 mM fresh DTT at pH 6.8) to remove residual glycerol, ATP and BDM in the fiber. Next, the fibers were incubated in rigor buffer containing 250µM mant-ATP for 5 min. After capturing background signal for initial 60 seconds, images were continuously taken while the fiber was washed with rigor buffer containing 4mM ATP to flush mant-ATP for next 5 minutes. All images were taken by Hamamatsu 1394 ORCA-ERA camera at 20x objective power under DAPI filter (excitation 360nm, emission 460nm) equipped on a Leica DMi8 Widefield Fluorescence microscope every 5 seconds at room temperature 23°C. Fluorescence intensity was measured in 2∼3 different regions of fiber (50 μm x 50 μm) plus one background region. Individual fiber fluorescence intensity of each time point was subtracted by background intensity and normalized by the averaged baseline value prior to mant-ATP washout. To measure single myosin nucleotide (ATP) turnover, biphasic pulse-chase method was used as previously described. (42). All the results were fit to the double exponential decay to obtain 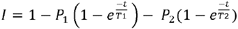, where I is the fluorescence intensity at any given time (t). P1 is defined as the relative proportions of fluorescence in the first fast phase exponent and P2 is defined as the relative proportions of fluorescence in the second slow phase exponent. T1 determines the lifetime of the first fast phase exponent and T2 determines the lifetime of the second slow phase exponent, which are the inverse of the rate of ATP turnover. The proportion of the SRX myosin is 2 × P2, and the percentage of the DRX myosin is 1 − (2 × P2).

### Loaded ATP turnover assays in permeabilized porcine single cardiomyocytes

The ATP turnover rate of myosin in permeabilized single cardiomyocytes (CM) under loaded conditions was conducted as described (25). Briefly, frozen porcine myocardium was cut into 10-15 mg pieces and permeabilized on ice in skinning solution (isolation buffer: 5.55 mM Na_2_ATP, 7.11 M MgCl_2_, 2 mM EGTA, 108.01 mM KCl, 8.91 KOH, 10 mM Imidazol, 10 mM DTT + 0.3% TritonX-100) with protease inhibitor cocktail (Sigma Aldrich) and phosphatase inhibitors (PhosSTOP, Roche). The tissue was homogenized with low-speed pulverization, skinned for 20 minutes at 4°C, and washed with isolation buffer to remove Triton. CMs were affixed to a force and length transducer using an ultraviolet-activated adhesive (Norland Optical Adhesives) and the sarcomere length was set to 2.1 μm. The CM was then washed in rigor buffer (6.41 mM MgCl_2_, 10 mM EGTA, 100 mM BES, 10 mM CrP, 50.25 mM Kpropionate, protease inhibitor, 10 mM DTT) to remove ATP and subsequently incubated in relaxing buffer made with 25 μM 2’-/3’-O-(N’-Methylanthraniloyl) adenosine-5’-O-triphosphate (mant-ATP, Enzo Life Sciences, Axxora LLC, Framingham, NY). CMs were then moved to relaxing buffer and fluorescence intensity acquired (excitation 352-402 nm, emission 417-444 nm; Horiba / PTI 814 Photomultiplier Detection System) continuously at 100 Hz for 1000 s at room temperature 23°C. The fluorescence signal was filtered using a second order Savitzky-Golay filter and normalized and fit to a double exponential function as described above. Following acquisition of the fluorescence decay curve, CMs were incubated in 1 μM OM and 7 μM piperine in relaxing solution for 10 minutes, and the assay repeated post-exposure, with the chase performed in the presence of drug. All analysis was performed using custom routines written in Matlab (Mathworks, 2020).

## Acknowledgments

This research used resources of the Advanced Photon Source, a U.S. Department of Energy (DOE) Office of Science User Facility operated for the DOE Office of Science by Argonne National Laboratory under Contract No. DE-AC02-06CH11357. This project is supported by grant P30 GM138395 from the National Institute of General Medical Sciences of the National Institutes of Health. The content is solely the authors’ responsibility and does not necessarily reflect the official views of the National Institute of General Medical Sciences or the National Institutes of Health. Predoctoral Fellowship Grants: AHA 23PRE1026275 and F31 HL168850 (VPJ);

## Author contributions

W.M, designed the experiments; V.J, T.S, W.M, and H.G performed the experiments; V.J, T.S, W.M, and C.G analyzed the data; W.M, S.S, D.K, and T.C.I. wrote the manuscript. All authors approved the final version of the manuscript.

## Competing interests

W.M and T.C.I. consults for Edgewise Therapeutics, but this activity has no relation to the current work.

## Data Availability

The datasets generated or analyzed during this study are included in this article. The raw data are available from the corresponding authors (Weikang Ma) on reasonable request.

